# Dual tRNA mimicry in the Cricket Paralysis Virus IRES uncovers an unexpected similarity with the Hepatitis C Virus IRES

**DOI:** 10.1101/230615

**Authors:** Vera P. Pisareva, Andrey. V. Pisarev, Israel S. Fernández

## Abstract

Co-opting the cellular machinery for protein production is a compulsory requirement for viruses. The Cricket Paralysis virus employs an <u>I</u>nternal <u>R</u>ibosomal <u>E</u>ntry <u>S</u>ite (IRES) to express its structural genes in the late stage of infection. Ribosome hijacking is achieved by a sophisticated use of molecular mimicry to tRNA and mRNA, employed to manipulate intrinsically dynamic components of the ribosome. Binding and translocation through the ribosome is required for this IRES to initiate translation. We report two structures, solved by single particle electron cryomicroscopy (cryoEM), of a double translocated CrPV IRES with aminoacyl-tRNA in the peptidyl site (P site) of the ribosome. CrPV IRES adopts a previously unseen conformation, mimicking the acceptor stem of a canonical E site tRNA. The structures suggest a mechanism for the positioning of the first aminoacyl-tRNA shared with the distantly related Hepatitis C Virus IRES.

---

Translation initiation is the most complex and highly regulated step of protein synthesis[1]. Canonical initiation results in the formation of an elongation-competent ribosome with an aminoacyl-tRNA base paired with messenger RNA (mRNA) at the peptidyl site (P site) of the ribosome. Viruses can hijack host ribosomes to produce viral proteins[2, 3]. A common strategy used by different types of viruses relies on structured RNA sequences at the ends of their mRNAs[4].These Internal Ribosomal Entry Sites (IRES) form specific three-dimensional structures and are able to manipulate and co-opt the host translational machinery[5].

IRES sequences are classified according to the subset of factors they require for initiation[4]. Type IV IRES sequences, including the Cricket Paralysis Virus IRES (CrPV IRES) and the Taura Syndrome Virus IRES (TSV IRES) do not require initiation factors and are the best studied IRESs. Biochemical and structural studies have provided a detailed view on how these approximate 200 nucleotide long sequences interact with and manipulate the ribosome[6, 7, 8].

A modular architecture of three pseudoknots (PKI, PKII and PKIII, **Fig. 1, A**) is crucial for these IRESs to establish a balance between structural flexibility and rigidity, essential for interaction with the ribosome and with two elongation factors (eEF2 and eEF1A) required for IRES translocation through the ribosome[9]. PKI mimics an anti-codon stem loop (ASL) of a tRNA interacting with its cognate mRNA codon, and plays an essential role in setting up the correct reading frame in the aminoacyl site (A site) of the ribosome[10].

**Fig.1.**
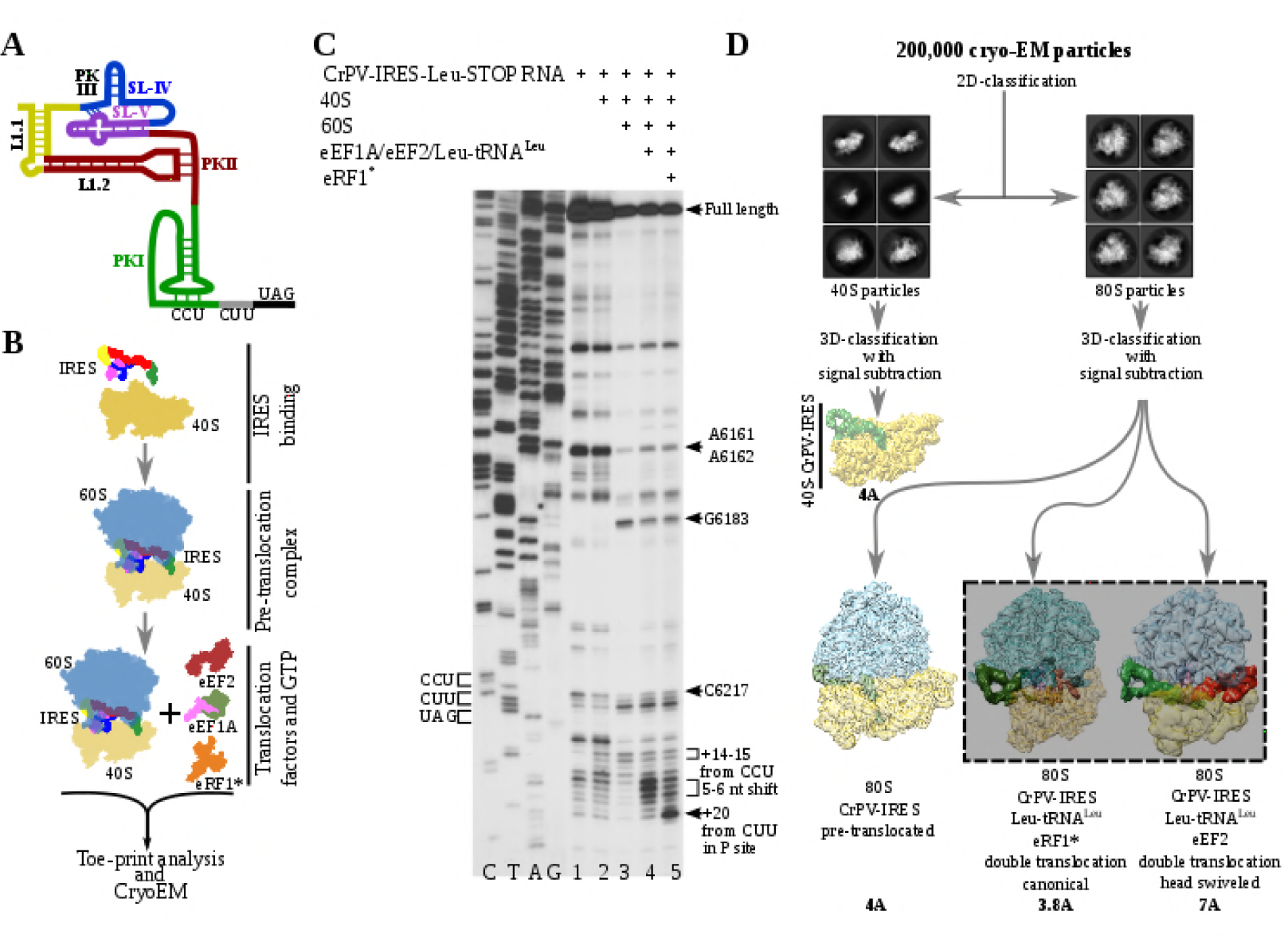
Experimental set up and cryoEM image processing workflow. (**A**) Secondary structure scheme of CPV IRES highlighting the modular architecture consisting of three pseudoknots (PKI, PKII and PKIII). (**B**) Diagram showing the *in vitro* reaction set up with purified components used for the toe-printing assays as well as for cryoEM. (**C**) Toe-print analysis of ribosomal complexes assembled with components indicated on top. Toe-prints corresponding to the pre-translocated complex are labeled as “+14-15 nt from CCU”. Toe-prints corresponding to the double translocated complex without and with eRF1* are marked as“5-6nt shift” and “+20 nt from CUU” respetively. Additional toe-prints previously attributed to different CrPV IRES/ribosome contacts are seen, in agreement with previous reports[15, 20]. (**D**) CryoEM data processing workflow employed to resolve the high compositional and conformational heterogeneity of the *in vitro* reconstituted complexes described in **B**. Four populations were resolved and refined to high resolution, including two exhibiting clear density for a double translocated CrPV IRES (squared).

A ribosome primed with a type IV IRES alternates between rotated and non-rotated configurations of the small ribosomal subunit, resembling a pretranslocation state of the ribosome with tRNAs immediately after peptidyl transfer[11, 12]. The rotated configuration is the substrate for eEF2 which, in its GTP-bound form, facilitates the translocation of the PKI into the P site, vacating the A site for the delivery of the first aminoacyl-tRNA[13, 14]. A second translocation event should take place for the first aminoacyl-tRNA to enter the P site, ending the unusual initiation pathway followed by this type of IRESs.

We report the visualization by means of single-particle electron cryomi-croscopy (cryoEM) of two related states of the mammalian ribosome with a double translocated CrPV IRES and P site aminoacyl-tRNA at 3.8 and 7 Ångstroms resolution. The head swiveling of the of the small ribosomal subunit plays a fundamental role in the late step of this translocation event, inducing a remarkable conformational change on the PKI of the CrPV IRES, which becomes disassembled, to mimic the acceptor stem of a E site tRNA.

To analyze the integrity and stability of complexes for cryoEM, we assembled ribosomal complexes with a double translocated CrPV IRES in a mammalian reconstituted system from individual components in the presence of GTP(**Fig. 1, B** and **C**). The translocation efficiency was monitored by toe-printing. Pre-translocation complex results in a +14-15 nt toe-print signal from CCU indicating the presence of PKI in the ribosomal A site (**Fig. 1, C**, lane 3). The addition of elongation factors leads to a 5-6 nucleotides toe-print shift showing the double translocation event (**Fig. 1, C**, lane 4). However, the similar intensity of several bands, with difference on 1 nucleotide, suggests instability of the double translocated IRES or frame ambiguity. To stabilize a double translocated complex, we introduced a UAG stop codon following the CUU coding triplet (**Fig. 1, A**). The supplementation of the reaction with a mutated and catalytically inactive version of the release factor 1 (eRF1*, AGQ mutation) causes a +20 nt toe-print signal from the CUU codon in the P site, which is in a good agreement with previous reports[15], indicating proper binding of eRF1* (**Fig. 1, C**, lane 5). The simultaneous decrease on intensity for the 5-6 nucleotides toe-print suggests a more homogeneous complex, suitable for structural studies.

Maximum likelihood particle sorting methods implemented in RELION[16] were applied to a large cryoEM dataset at two different stages (**Fig. 1, D**). An initial classification in two dimensions allowed for the separation of full ribosome (80S) from small subunit (40S) particles. The two sorted subgroups were further classified using masking methods with signal subtraction with focus in the inter-subunit space, where this type of IRES binds as well as canonical translation factors. The L1 stalk was also included in the masked area to allow for a wider sampling. This strategy revealed, in a single classification step, several sub-populations, reflecting the heterogeneity of the sample. A binary 80S/CrPV IRES complex in a pre-translocation conformation could be identified as well as two sub-populations with CrPV IRES in a double translocated state. In the double translocated reconstructions, the ribosome adopts a non-rotated configuration of the small subunit, with clear density for aminoacyl-tRNA in the P site and a either eEF2 or eRF1* in the A site (**Fig. 1, D** and **Fig. 2 and 3**).

**Fig.2.**
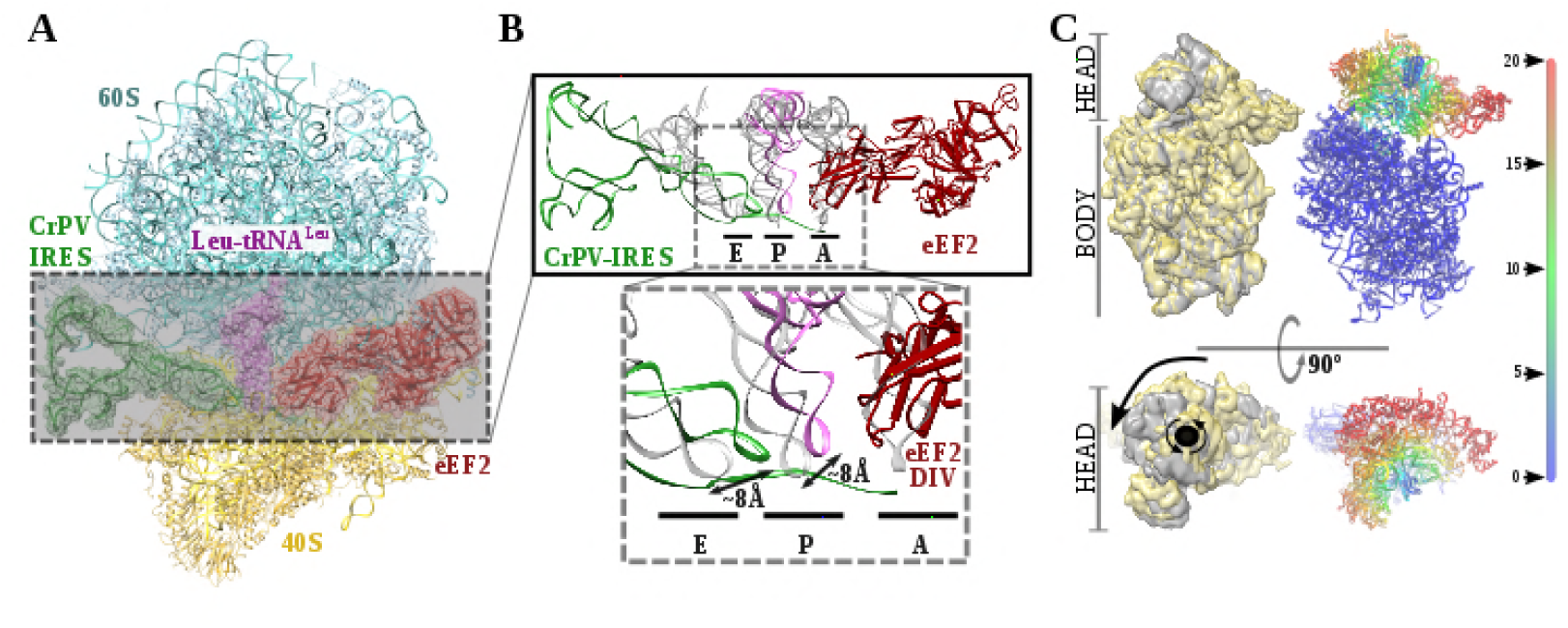
Structure of a double translocated CrPV IRES intermediate with eEF2. (**A**) Overview of a mammalian ribosome with double translocated CrPV IRES (green), aminoacyl-tRNA (purple) and eEF2 (red). (**B**) Top, detailed view of the ribosomal sites E, P and A in the structure with a double translocated CrPV IRES with eEF2. Canonical tRNAs (from PDB ID 4V5C) are depicted as semi-transparent grey cartoons. Bottom, zoomed view highlighting the displacement of both aminoacyl-tRNA and CrPV IRES PKI from canonical positions. Domain IV of eEF2 occupies the A site. (**C**) On the left, two views of the experimental density for the 40S of the two double translocated reconstructions where it can be appreciated the swiveled configuration of the 40S head in the complex with eEF2 (grey). For comparative purposes, the map for the eRF1* containing complex (yellow) has been low pass filtered to a similar resolution as the eEF2 containing complex (grey). Right, atomic refined model colored according to the root-mean squared displacement (RMSD) with red and blue indicating the highest and lowest values, respectively (in Ångstroms).

**Fig.3.**
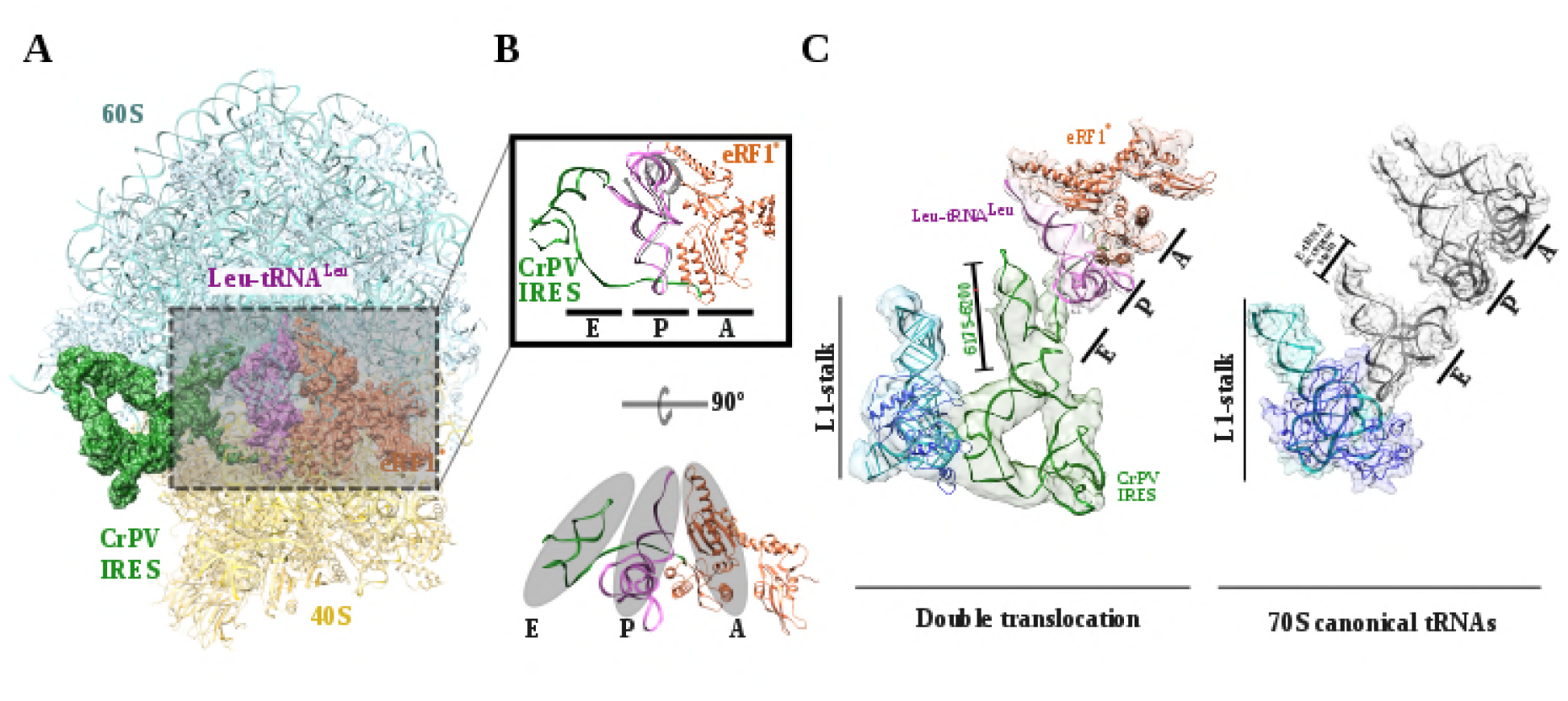
Structure of a double translocated CrPV IRES. (**A**) Overview of a mammalian ribosome with double translocated CrPV IRES (green), aminoacyl-tRNA (purple) and eRF1* (orange). (**B**) Top, detailed view of the ribosomal E, P and A sites with a canonical configuration for the aminoacyl-tRNA in the P site, eRF1* in the A site and a disassembled CrPV IRES PK I in the E site (green). Canonical P site tRNA (from PDB ID 4V5C) is depicted as semi-transparent grey cartoon. Bottom, a 90 degrees rotated view showing the ASL-like part of the CrPV IRES PKI after disassembling populating the E site of the 60S, in a similar position as the acceptor stem of a canonical E site tRNA. (**C**). The position of the three ligands in the double translocated complex with eRF1* in comparison with canonical tRNAs. The L1 stalk is depicted in cyan.

Recent studies by single-molecule FRET (smFRET) have characterized the kinetics of the translocation events required for the CrPV IRES to deliver an aminoacyl-tRNA to the P site of the ribosome[8, 17]. Slow movements of the CrPV IRES compared with canonical translocation of tRNAs are evident from this data and explain the capturing of a late-stage intermediate of translocation with eEF2 in our dataset (**Fig. 2, A-C**). The conformation adopted by CrPV IRES in this intermediate state is similar to the conformation reported for the single translocated state[15], with the SL-IV and SL-V detached from the 40S and exposed to the solvent. PKI is in an intermediate position between the P and E sites of the small subunit as well as the aminoacyl-tRNA is in an intermediate position between the A and P site of the 40S (**Fig. 2, B**). Domain IV of eEF2 (the eukaryotic ortholog of bacterial EF-G) occupies the A site. This configuration is maintained by a distinctive swiveled configuration of the 40S head, resembling one of the late stages recently reported for the first translocation event (**Fig. 2, C**)[14].

The most populated class of particles represent a double translocated CrPV IRES with aminoacyl-tRNA in a canonical configuration in the P site and eRF1* in the A site (**Fig. 3**). Both aminoacyl-tRNA and eRF1* populate conformations recently described, with the characteristic bent of the mRNA at the stop codon[18]. The small subunit in this reconstruction is in a non-rotated configuration and the 40S head is not tilted or swiveled (**Fig. 3**). CrPV IRES has undergone a conformational change that mainly affects PKI, but also the relative orientation of PKII and PKIII. Upon back-swiveling of the 40S following eEF2 departure, the aminoacyl-tRNA is placed in its final canonical position in the P site (**Fig. 3, B**). This event triggers the disassembly of the CrPV IRES PKI. Although the mRNA-like part remains placed in the E site of the 40S, the ASL-like segment experiences a pronounced displacement to occupy the E site of the 60S, now mimicking the acceptor stem of a canonical E site tRNA (**Fig. 3, B** and **C**). The L1.1 part of CrPV-IRES remains attached to the L1 stalk along this process whose displacement relative to the 60S is similar to the one described in the complex with eEF2 and a non-hydrolyzable GTP analog or after the first translocation (**Fig. 4. A**)[13, 15]. The back-swiveling of the 40S head upon eEF2 departure is also involved in a new relative orientation of PKII and PKIII (**Fig.3, B** and **C**). In the swiveled configuration (**Fig. 4, B** left), SL-IV and SL-V, components of PKII, are exposed to the solvent, in a similar position described for the single translocated CrPV IRES[15]. The eukaryotic specific protein eS25, a key element of the small subunit involved in early recruitment of the CrPV IRES as well as in the positioning of the IRES in the pre-translocation stage[19], is not interacting with the IRES (**Fig. 4, B**, left arrow). Upon back swiveling of the 40S head, a new interaction is established between the CrPV IRES and eS25 (**Fig.4, B** right, arrow) not involving SLV like in the pre-translocated complex. The new interaction stabilizes the CrPV IRES in a distinctive conformation, with PKII and PKIII assembled but with a wider relative orientation (**Fig. 4, C**) and the PKI disassembled with residues 6175 to 6200, corresponding to the ASL mimicking part of PKI, populating a space corresponding to the acceptor stem of a canonical E site tRNA.

**Fig.4.**
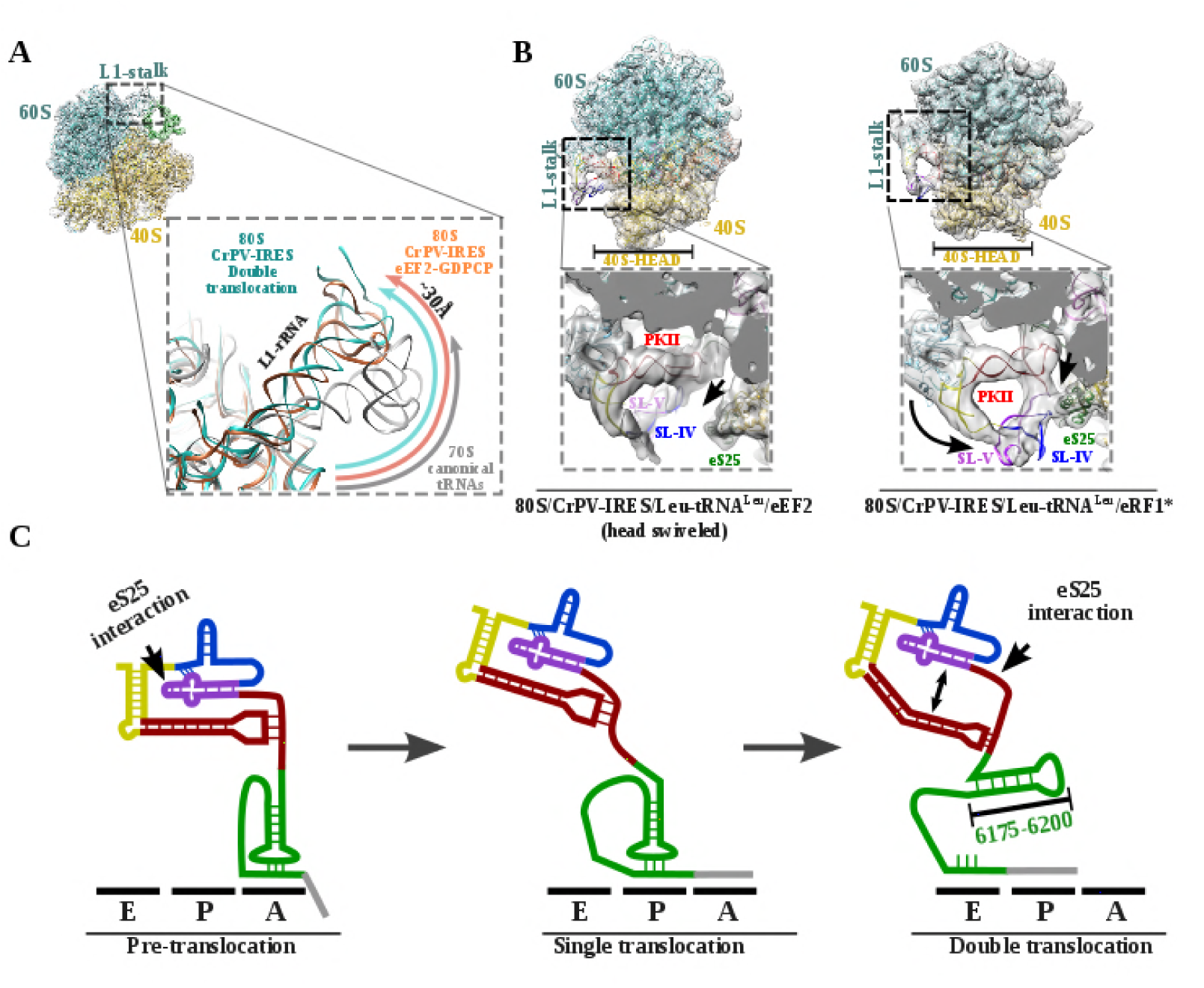
L1 stalk position and conformational change on double translocated CrPV IRES. (**A**) Top left, overview of the double translocated ribosome complex with eRF1* with the L1 stalk region highlighted. Main view, L1 stalk in the double translocated complex (cyan) is displaced from the position acquired in a complex with canonical tRNAs (grey) with a magnitude of approximately 30Å. This displacement is similar to the one reported for the pre-translocated complex with eEF2 and a non-hydrolyzable GTP analog (orange)[13]. (**B**) Conformational transition observed in CrPV IRES upon back-swiveling of the 40S head. Left, due to a swiveled 40S head configuration in the double translocated complex with eEF2, SL-IV and SL-V remain solvent exposed, as in the single translocated complex[15], and detached from the ribosomal protein eS25 (green). Right, once the head of the 40S relocates to its nonswiveled position, CrPV IRES acquires a new conformation involving a new interaction with eS25 (green). (**C**) Scheme showing the secondary structure of CrPV IRES in the pre-translocated state (left), after a single translocation (center) and after the double translocation (right). Arrows indicate the repositioning of PKII and PKIII as well as the new interaction established with ribosomal protein eS25.

The structures provide a structural view of the archetypical CrPV IRES in the final stage of initiation, after transitioning through the ribosome. Combining the structure with published biochemical and smFRET data allows us to propose a comprehensive working model for how the CrPV IRES (and type IV IRES in general) recruits, manipulates and redirects host ribosomes for the synthesis of its own proteins (**Fig. 5**). As suggested by classic crosslinking experiments[20], smFRET data[8] and cryoEM reconstructions[7, 13], CrPV IRES initially assembles a binary 80S/CrPV IRES complex by either directly recruiting empty 80S or by a step-wise pathway in which CrPV IRES first recruits the 40S subunit and then the 60S subunit (**Fig. 5**, bottom left). Once the binary 80S/CrPV IRES is assembled, the 40S oscillates between rotated and non-rotated states, with PKI inserted in the A site and minimum changes in the overall conformation of the IRES[11, 12]. These movements are coupled to oscillations of the L1 stalk. The rotated state is the substrate of eEF2, which, in its GTP-bound form, induces an additional rotation of the small subunit and additional displacement of the L1 stalk, to facilitate the translocation of the PKI from the A to the P site (**Fig. 5, A** top left). Back rotation and back swiveling of the 40S, combined with ribosome-induced GTP hydrolysis by eEF2 results in the first translocation event of the CrPV IRES, positioning PKI in the P site, mimicking a translocated, canonical aminoacyl-tRNA. This intermediate is unstable and prone to back-translocation[15], unless a cognate aminoacyl-tRNA, delivered to the ribosome in complex with eEF1A and GTP, captures the frame in the A site of the ribosome[8]. Formation of this complex is a rate-limiting step in this kinetically driven process[8]. In the single translocated IRES state, SL-IV and SL-V, which are initially attached to the ribosome, are solvent exposed, the PKI occupy the P site of the 40S and an aminoacyl-tRNA occupies the A site. It is reasonable to assume this state will oscillate between rotated and non-rotated configurations of the small subunit as a canonical pre-translocation complex with tRNAs[21]. The second translocation step is required to place the first aminoacyl-tRNA in the P site and thus finish the initiation phase of translation (**Fig. 5, A** bottom right). At this stage, CrPV IRES translocation should be coupled with the movement of the aminoacyl-tRNA occupying the A site and this seems to happen with a conformation of the IRES similar to the one reported for the first translocation[15]. This conformation is maintained until the very last moment as the intermediate captured here with eEF2 presents a back-rotated configuration of the 40S[14]. However, a pronounced swiveling of the 40S head is in place, probably induced by the presence of eEF2[14]. Once eEF2 leaves, the back-swiveling movement of the 40S head triggers a dramatic conformational change in the CrPV IRES: PKI is disassembled resulting in the ASL-like segment relocating to mimic the acceptor stem of a canonical E site tRNA. The mRNA-like element of the disassembled PKI remains in the E site of the 40S. These conformational changes in the PKI of the CrPV IRES upon back swiveling are combined with a reconfiguration of the relative positioning of PKII and PKIII. This new conformation is stabilized by a newly reported IRES/40S interaction with the ribosomal protein eS25, which is also involved in the early recruitment of the IRES to the 40S[13].

**Fig.5.**
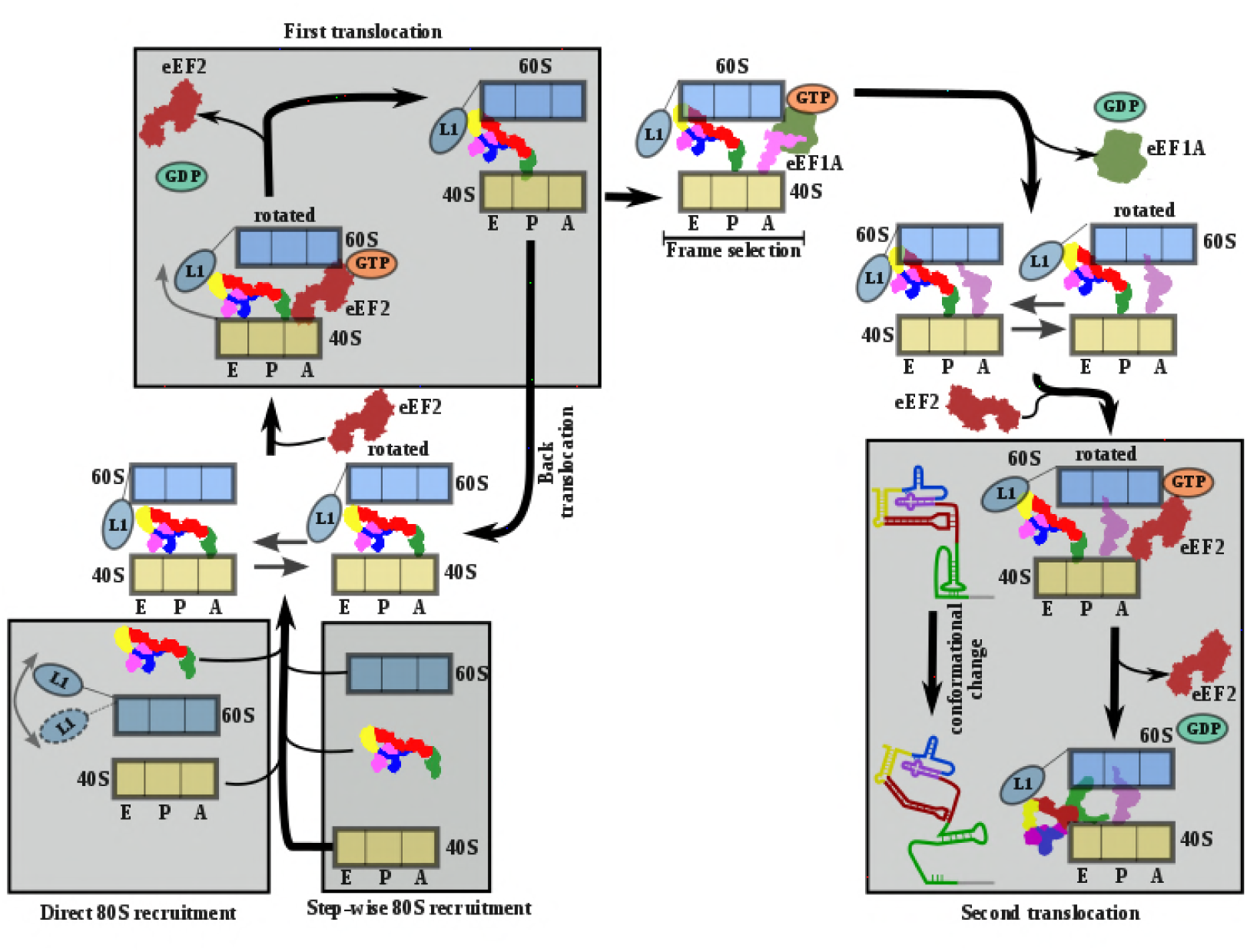
Comprehensive model describing CrPV IRES strategy to hijack host ribosomes. Bottom left, CrPV IRES can directly recruit 80S to assemble a binary 80S/CrPV IRES complex which, in its pre-translocation state, oscillates between rotated and non-rotated configurations of the 40S. However, a step-wise pre-translocation complex formation involving an initial interaction with the 40S, followed by recruitment of 60S, is more efficient and is favored. Top left, first translocation event involving the displacement of CrPV IRES PKI from the A site in order for the first aminoacyl-tRNA to be delivered as a ternary complex with eEF1A and GTP. In the absence of an A site ligand this state is unstable and prone to back translocation[15]. According to smFRET studies, the frame is not defined until the first condon/anticodon interaction is established[8]. Top right, presumably a single translocated complex with A site aminoacyl-tRNA alternates between rotated and non-rotated configurations of the 40S as a *bona fide* pre-translocation complex with two tRNAs. Bottom right, binding of eEF2 in its GTP form assists in the translocation of CrPV IRES and the first aminoacyl-tRNA which is achieved through a conformational change in the CrPV IRES involving the disassembling of the PKI and reorientation of PKII and PKIII.

The conformational change described here for the CrPV IRES following translocation through the ribosome, unexpectedly resembles the transition observed for the Hepatitis C Virus (HCV) IRES upon aminoacyl-tRNA delivery to the P site (**Fig. 6**)[22, 23]. The HCV IRES belongs to a different class of IRES, due to it requires some canonical factors to initiate translation[4, 5]. It also interacts with the ribosome in a different manner[24]. However, a large stem (**Fig. 6**, domain II, blue) reaches the E site of the 40S and is maintained base paired with the mRNA-like part of this IRES by a tilted configuration of the 40S head (**Fig. 6, B**)[23]. Upon delivery of initiator tRNA to the P site, the head recovers its non-tilted configuration resulting in the repositioning of the domain II to occupy a similar space as the CrPV IRES in the E site of the 60S (**Fig. 6**, right).

**Fig.6.**
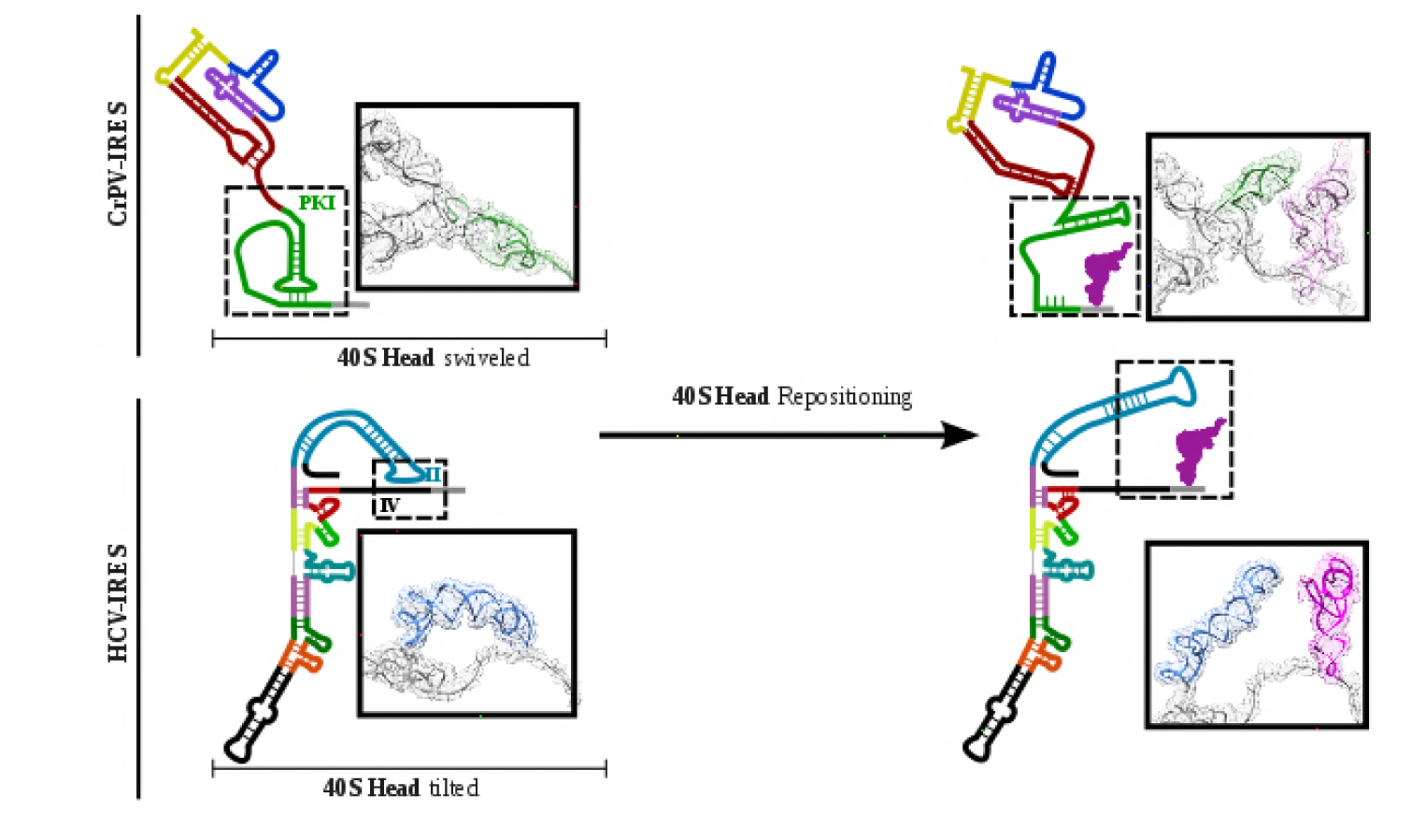
CrPV IRES and HCV IRES experiments a similar structural transition upon first aminoacyl-tRNA delivery to the ribosomal P site. The structural transition experienced by the CrPV IRES upon delivery of the first aminoacyl-tRNA to the P site (top) is similar to the one described for the HCV IRES (bottom). CrPV IRES PKI remains assembled and in the vicinity of the E site of the 40S due to a swiveled configuration of the 40S head. The HCV IRES maintains a similar internal interaction of domain II by a tilted configuration of the 40S head. In case of both IRESs, a reconfiguration involving back-positioning of the 40S head plus the placement of a structural element of the IRES in the vicinity of the 60S E site, facilitates the delivery of the first aminoacyl-tRNA to the P site of the ribosome, finalizing the initiation stage of translation.

Therefore, to assemble translationally competent ribosomes, distantly related IRESs have converged on a similar mechanism to regulate the placement of the first aminoacyl-tRNA in the P site of the ribosome, by resembling endogenous tRNA states.

## Acknowledgements

We acknowledge Bob Grassucci for technical assistance in data acquisition, Harry Kao for computing and Alan Brown for critical reading of the manuscript. CryoEM data was acquired at the Simons Electron Microscopy Center in the New York Structural Biology Center. We are grateful to Ed Eng, Bill Rice, Laura Kim, Anchi Chen, Clint Potter and Bridget Carragher for support at all stages of this project. Density maps have been deposited at the EMDB with accession codes XXXX and YYYY. Atomic coordinates have been deposited in the PDB with accession codes xxxx and yyyy. V.P.P. and A.V.P. are funded by National Institutes of Health [GM097014 to A.V.P.] grant.

## METHODS

### Plasmids

Expression vector for His-tagged eRF1*(AGQ mutant)[1] and transcription vector for Leu-tRNA have been previously described[2]. Transcription vector for CrPV-Leu-STOP was constructed inserting a T7 promoter sequence upstream of CrPV IGR IRES sequence followed by the two first coding triplets and an EcoRI site, using pUC19 as a scaffold vector. Site-directed mutagenesis was employed to change the first coding triplet to CUU encoding leucine and the second coding triplet to a stop (UAG) codon, rendering the CrPV-Leu-STOP construct. CrPV-Leu-STOP RNA and Leu-tRNA were transcribed using T7 RNA polymerase.

### Purification of translation components and aminoacylation of Leu-tRNA

Native 40S and 60S subunits, eEF2, rabbit aminoacyl-tRNA synthetases[3], and eEF1A[4]were prepared as previously described. Recombinant eRF1* was purified according to a previously described protocol[3]. *In vitro* transcribed Leu-tRNA was aminoacylated with leucine in the presence of rabbit aminoacyl-tRNA synthetases as previously described[2].

### Assembly of ribosomal complexes

To reconstitute different ribosomal complexes, we incubated 1.8 pmol 40S ribosomal subunits with 2 pmol CrPV-Leu-STOP RNA in a 20 μl reaction mixture containing buffer A (20 mM Tris-HCl, pH 7.5, 100 mM KCl, 2.5 mM MgCl_2_, 0.1 mM EDTA, 1 mM DTT) with 0.4 mM GTP for 5 min at 37°C. Then, the reaction mixture was supplemented with 2.5 pmol 60S ribosomal subunits and additionally incubated for 5 min at 37°C. Next, we added 10 pmol eEF1A, 3pmol eEF2, and 0.4 pg Leu-tRNA^Leu^, and incubated for 5 min at 37 ^°^C. Finally, ribosomal complexes were incubated with 20 pmol eRF1 (AGQ) for 5 min at 37°C. We analyzed the assembled ribosomal complexes via a toe-printing assay essentially as described[5].

### CryoEM sample preparation and data acquisition

Aliquots of 3μl of assembled ribosomal complexes at concentration of 80100 nM were incubated for 30 seconds on glow-discharged holey gold grids (UltrAuFoil R1.2/1.3[6]), on which a home-made continuous carbon film (estimated to be 50Åthick) had previously been deposited. Grids were blotted for 2.5s and flash cooled in liquid ethane using an FEI Vitrobot. Grids were transferred to an FEI Titan Krios microscope equipped with an energy filter (slits aperture 20eV) and a Gatan K2 detector operated at 300 kV. Data was recorded in counting mode at a magnification of 130,000 corresponding to a calibrated pixel size of 1.08 Å. Defocus values ranged from 1.6-3.6 μm. Images were recorded in automatic mode using the Leginon[7] software and frames were aligned with Motioncor2[8] and checked on the fly using APPION[9].

### Image processing and structure determination

Contrast transfer function parameters were estimated using GCTF[10] and particle picking was performed using GAUTOMACH without the use of templates and with a diameter value of 260 pixels. All 2D and 3D classifications and refinements were performed using RELION[11]. An initial 2D classification with a 4 times binned dataset identified all ribosome particles. A second 2D classification step with 2 times binned data was employed to separate 80S from 40S particles. A consensus reconstruction with all 80S particles was computed using the AutoRefine tool of RELION whose resulting map was use to built a mask containing the inter-subunit space and the L1 stalk. 3D classification with signal subtraction using the previously described mask and a T value of 10 allowed for the identification of several population of ligands inside the mask, namely empty ribosomes, pre-translocated CrPV IRES and double translocated populations with aminoacyl-tRNA and eEF2 or eRF1*. Final refinements with unbinned data for the classes selected yielded high resolution maps with density features in agreement with the reported resolution. Local resolution was computed with RESMAP[12].

### Model building and refinement

Models for the mammalian ribosome, Leu-tRNA^Leu^, eEF2 and eRF1* were docked into the maps using CHIMERA[13] and COOT[14] was used to manually adjust the L1 stalk and rebuild CrPV IRES using our previous model as initial step. An initial round of refinement was performed in Phenix using real space refinement with secondary structure restrains[15]. A final step of reciprocal-space refinement using REFMAC was performed[16]. The fit of the model to the map density was quantified using FSCaverage and Cref.

## Supplementary figure legends

**Fig.S1.**
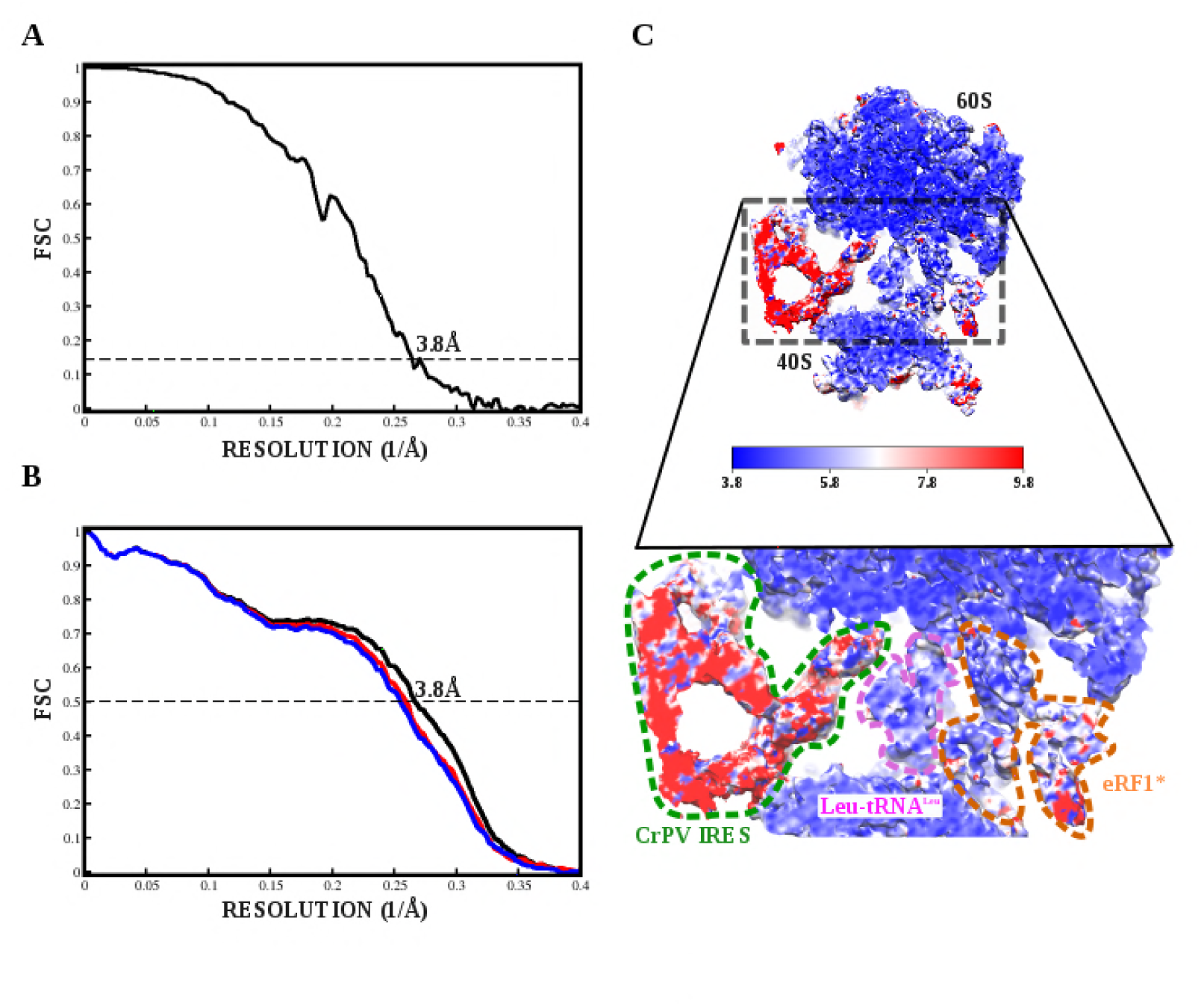
Fourier Shell Correlation curves and local resolution estimation for the 80S/CrPV IRES/Leu-tRNA^Leu^/eRF1* complex. (**A**) Fourier Shell Correlation (FSC) computed for the two half maps of the final subset of particles after classification. The resolution is estimated to be 3.8Å using the 0.143 criterion. (**B**) Map-versus-model cross validation FSC. The final model was validated using standard procedures. FSC of the refined model against half map 1 (blue) overlaps with the FSC against half map 2 (red, not included in the refinement). The black curve corresponds to the FSC of the final model against the final map (**C**) Slice through the final, unsharpened map colored according to the local resolution as reported by RESMAP.

**Fig.S2.**
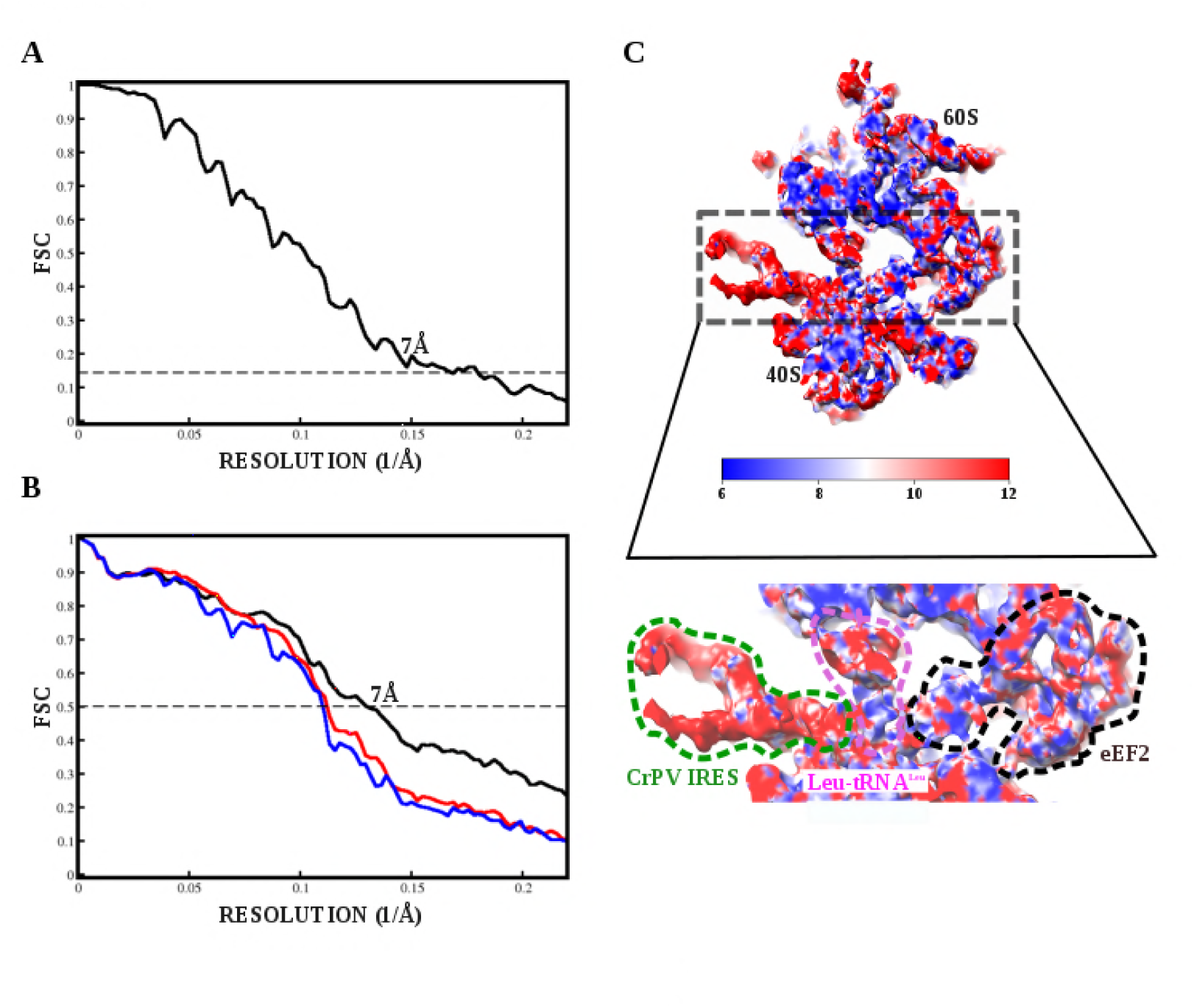
Fourier Shell Correlation curves and local resolution estimation for the 80S/CrPV IRES/Leu-tRNA^Leu^/eEF2 complex. (**A**) ourier Shell Correlation (FSC) computed for the two half maps of the final subset of particles after classification. The resolution is estimated to be 7Å using the 0.143 criterion.(**B**) Map-versus-model cross validation FSC. The final model was validated using standard procedures. FSC of the refined model against half map 1 (blue) overlaps with the FSC against half map 2 (red, not included in the refinement). The black curve corresponds to the FSC of the final model against the final map (**C**) Slice through the final, unsharpened map colored according to the local resolution as reported by RESMAP.

